# The implicit influence of pitch contours and emotional timbre on P300 components in an own-name oddball paradigm

**DOI:** 10.1101/2023.11.30.569381

**Authors:** Estelle Pruvost-Robieux, Coralie Joucla, Sarah Benghanem, Rudradeep Guha, Marco Liuni, Martine Gavaret, Jean-Julien Aucouturier

**Affiliations:** Neurophysiology and Epileptology Department, GHU Paris Psychiatrie et Neurosciences, Sainte Anne Hospital, Paris, France; Université Paris Cité, Paris; INSERM UMR 1266, Institut Paris Neurosciences et Psychiatrie (IPNP), Paris; Science and Technology of Music and Sound Lab (IRCAM/CNRS/Sorbonne Université), Paris, France; Université de Franche-Comté, SUPMICROTECH, CNRS, institut FEMTO-ST, F-25000 Besançon, France; Medical ICU, Cochin Hospital, Assistance Publique - Hôpitaux de Paris (AP-HP), 27 Rue du Faubourg Saint-Jacques, 75014, Paris, France; Product designer, Mezzo Forte, Paris, France

**Keywords:** pitch perception, timbre perception, evoked potentials, own-name, oddball

## Abstract

**Objective:** Previous evidence has suggested that paralinguistic features of speech stimuli may influence the characteristics of P300 components when used in clinical evaluation of consciousness. However, it remains unknown what exact acoustic components of speech are influential in such tasks, and whether they are capable of interacting with attentional deployment (as indexed by P300) in an implicit manner.

**Methods:** To study this question, we adapted here an auditory oddball-paradigm used in clinical practice (the “own-name” paradigm), and tested whether systematic transformations of pitch contours (i.e. rising or falling intonation) and emotional timbre (i.e. smiling or rough voices) on the participants’ names influenced P300 responses in 24 healthy participants.

**Results:** P300 responses to rising pitch contours were smaller than to falling pitch contours, possibly reflecting an interference of the rising contours with participant attention in the deviant-counting task. No such difference was observed with emotional timbre variations.

**Conclusion and Significance:** These results suggest that the cognitive resources involved in pitch contour processing overlap more strongly than timbre with the resources required to count own-name deviants, and that rising pitch contours should be tested prospectively as a way to increase the saliency of consciousness evaluation in unconscious patients.

## 1. Introduction

The expressive content of a voice is an important cue in human communication (Van Lancker Sidtis, 2018). Expressive intonations may reflect emotions, like happiness or sadness (Juslin & Laukka, 2003); epistemic attitudes, like doubt or certainty (Goupil et al., 2021) or speaker behaviors, such as trustworthiness or dominance (Ponsot et al., 2018; Wichmann, 2000).

Among all the acoustic characteristics of speech signals, certain prosodic dimensions are particularly relevant for such perceptions. The mean pitch (or fundamental frequency) of voices (i.e. whether a voice is lower or higher than usual) has for instance been associated with positive and/or aroused emotional states (Bachorowski & Owren, 1995) and social dominance/submissiveness (Mitchell & Ross, 2013). Beyond their mean level, pitch variations within an utterance also carry communicative meaning. Rising pitch contours, for instance, are described as more friendly and trustworthy (Ponsot et al., 2018; Torre et al., 2016), while falling pitch has been associated with assertiveness and increased speaker reliability (Goupil et al., 2021). In association to pitch, timbre (or voice quality) was also shown relevant to discriminate between attitudes and emotions (Grichkovtsova et al., 2012). For example, at similar pitch, the spectral features of smiling voices modulate facial mimicry in listeners (Arias et al., 2018); in laughter, whether the timbre is voiced (song-like) or unvoiced (grunts) modulates listeners’ positive responses (Bachorowski & Owren, 2001). Conversely, the coarse and rough aspect of certain voices is known to elicit aversion and recruit increased attention in listeners (Arnal et al., 2019).

Whereas the processing of vocal pitch and timbre is often investigated in explicit rating tasks (for a review, see Juslin et Laukka 2003), a number of behavioral studies also highlight that the implicit processing of these acoustic cues can modulate the attention of the participant. For instance, rough voices appear to increase synchronized activity in supra-temporal, limbic and frontal areas and this synchronization is associated with increased attention and aversion in listeners (Arnal et al., 2019). In the same spirit, the rising pitch of an utterance has been demonstrated to influence memory recall for pseudo-words, even when prosody is irrelevant to the task (Goupil et al., 2021). Finally, exposure to task-irrelevant environmental sounds (with or without emotional content) can modulate visual attention and facilitate visual search (Asutay & Västfjäll, 2017).

From a clinical point of view, documenting such implicit modulations of attention is crucial in non-responsive patients’ assessments. Demonstrating the implication of a disorder of consciousness (DOC) patient in a voluntary top-down attention task is an important signature of consciousness (Naccache, 2018). However, because such patients are unresponsive, practitioners cannot be sure that task instructions have really been understood or followed if a given patient fails to do a task. For this reason, even if a few active paradigms have been developed in DOC evaluation (Bekinschtein et al., 2009; Schnakers et al., 2008), most of clinical tools rely on passive paradigms, such as mismatch negativity (MMN) or P300 responses in auditory oddball tasks (André-Obadia et al., 2018). For instance, the “own-name paradigm”, where the patient’s own-name is embedded within more frequent tone bursts, is routinely performed in clinic practice (Fischer et al., 2008) to record a P300 response. Such passive tasks typically have good positive predictive value for awakening: patients who succeed in such tasks have a high probability of awakening, but little can be predicted in terms of subsequent cognitive disabilities (André-Obadia et al., 2018; Benghanem et al., 2022; Pruvost-Robieux, Marchi, et al., 2022). Moreover, these clinical tools typically have lower negative predictive value, as little can be predicted about patients who do not recruit attentional resources as expected. Knowing whether task-irrelevant cues, such as the prosodic expression of a participant’s own-name, modulate attentional recruitment in covert paradigms could provide clinical markers of higher-level cognitive function and, in fine, possibly represent a way to improve prognostication for DOC patients.

In a previous study, using a retrospective design, we tested the impact of the pitch contours used when recording own-name stimuli for testing N=251 DOC patients in two French Hospitals, and found that stimuli recorded with a rising-pitch intonation were statistically associated with shorter P300 latencies (Pruvost-Robieux, André-Obadia, et al., 2022). This result suggested that pitch modulation implicitly modifies patient’s results in the own-name paradigm, possibly by a modulation of attentional systems. However, because this study was performed with a retrospective design with many possible biases (patients with uncontrolled clinical conditions, etiologies of DOC, time spent since the onset of DOC, each only exposed to one type of prosody), no causal conclusions can be drawn about the general impact of prosody on P300 responses in patients or healthy controls. For these reasons, there is a need to investigate the relationship between modulations of own-name prosody and the resulting late auditory event related potentials (ERPs) in a prospective, within-subject study where subjects are evaluated with different prosodic versions of their own-name.

A recent EEG study focused on the role of timbre and pitch in evoking responses linked to the perception of the emotional content of a voice (Nussbaum et al., 2022). In this study, pitch was found relevant for EEG responses to happy, fearful and sad voices, while timbre impacted responses to voices expressing pleasure. However, the task was to explicitly judge the emotional content of stimuli, making prosody task-relevant. The question therefore remains whether pitch and timbre can implicitly modulate P300 indices of attention in a prosody-irrelevant task such as performed in ICU.

To do this, we subjected a group of N=24 healthy participants to a covert-attention EEG task adapted from the “own-name” paradigm routinely used in ICU to evaluate DOC patients. Sequences of standard tones and own-name deviants were presented. Participants were asked to covertly count the number of deviants, a task which is both known to generate clear P300 responses (Squires et al., 1976), and for which participants do not need to process the prosody of the sound. Without participants’ knowing, we digitally manipulated the own-name deviants either in pitch (rising or falling intonation) or emotional timbre (smiling or rough voices), using state-of-the-art voice transformation techniques (Bedoya et al., 2021), and looked whether these different prosodies implicitly influenced the resulting P300 characteristics.

## 2. Materials and methods

### 2.1 Study participants

Twenty-four healthy volunteers participated in the study (13 males; mean age = 25.1 years, SD 4.8, all right-handed). None of the participants reported auditory impairment or history of neurologic and psychiatric disorder. The experimental protocol was approved by Institut Européen d’Administration des Affaires (INSEAD)’s Institutional Review Board (protocol ID: 2021 – 51). Each participant provided an informed consent form before the beginning of the study. Participants were financially compensated for their time (20 euros / participant).

### 2.2 Audio recordings

We synthetized audio recordings of each participant’s first name using a commercial text-to-speech software (IBM Watson). Own-name stimuli duration depended on the duration of the participant first-name (in the limit of 1200ms) and was not altered by the acoustic transformations. To normalize the recordings according to timbre and pitch, we used the same male voice for synthesizing all stimuli, and then algorithmically flattened the pitch contour of the stimuli at the constant pitch value of 130 Hz using the CLEESE toolbox (Burred et al., 2019). We also normalized all stimuli in terms of RMS (Root-Mean-Square). We then used algorithmic manipulations to generate four variants of each own-name recording: two manipulated in pitch (‘rising’ and ‘falling’) and two in timbre (‘smile’ and ‘rough’). Examples of stimuli are available as supplemental material. Rising- and falling-pitch deviants were created using CLEESE, with which we forced the pitch contour of each stimuli to match the (roughly, linearly increasing or decreasing) contour found in our previous retrospective study (Pruvost-Robieux, André-Obadia, et al., 2022). The rising deviant had increasing pitch from 120 to 170 Hz. The falling deviant had decreasing pitch from 140 to 80 Hz.

Smiling and rough timbre deviants were created with two previously validated algorithms from our previous work: smiling deviants were created by transforming the spectral envelope of the sound using a phase vocoder technique, in a way that simulates the vocal tract transformations happening in smiled speech (Arias et al., 2020). Resulting speech stimuli were validated to be perceived more positive and arousing (Bedoya et al., 2021) and to trigger spontaneous facial imitations (Arias et al., 2018). Rough deviants were created by adding pitch-synchronous temporal modulations to the sound, mimicking the non-linearities involved in rough voices (Gentilucci et al., 2018). Resulting speech stimuli were validated to sound more negative and more aroused (Bedoya et al., 2021) and implicitly modulate auditory attention in spatial localization tasks (Ollivier et al., 2019). Finally, in all conditions, the standard sound associated to all type of deviants was a simple 75ms pure tone (1000Hz), generated with custom python code.

### 2.3 Oddball paradigms

Each participant took part in two successive oddball paradigms (one for pitch, one for timbre, duration about 15 min each), separated with a five-minute break. In the “pitch” paradigm, pure-tone standards were intermixed with the three pitch-deviant stimuli: rising, falling and flat (original flat) own-name. In the “timbre” paradigm, pure-tone standards were intermixed with the three timbre-deviant stimuli: smiling, rough and neutral. We randomly counterbalanced the order of the two oddball paradigms across participants. Each oddball paradigm included 1200 stimuli with 960 (80%) standards and 240 (20%) own-name deviants (6.6% of each deviant type). Each paradigm started by an uninterrupted sequence of 10 standard sounds, and sound presentation was then semi-randomized so that each deviant was followed by at least one standard sound. The inter-stimulus interval was set at 600ms for both standards and deviants, with a random jitter of +/- 5ms.

### 2.4 Procedure and EEG recording

Participants were comfortably settled in front of a screen fixation-cross and started to listen to the experiment. For the four first participants, stimulus presentation was controlled using the Presentation® software (neurobehavioral systems); for the remaining 20 participants, we used Python Psychopy (Peirce, 2009). Sounds were delivered bilaterally through earphones.

In both paradigms, participants were asked to covertly count the number of own-name deviants presented in the sequence, while ignoring standard sounds. Participants were explicitly instructed to count deviants regardless of their prosodic variations, which we described as arbitrary and task-irrelevant. Participants were unaware of the true purpose of the experiment, which was to study the influence of pitch and timbre modulations on their ability to ignore such prosodic variations while counting deviants.

Electroencephalographic (EEG) data were recorded using a 64-channel device (actiCHamp, Brain Products GmbH, Germany), with a sampling rate of 1000 Hz. Bandpass was set between 0.01-100Hz. EEG sensors were disposed according to the 10-10 system (Seeck et al., 2017). Cz was set as the reference electrode. Ground electrode was set on Fpz. Sound onset triggers were sent to the EEG acquisition computer by a Cedrus StimTracker (Cedrus Corporation, San Pedro, CA) to control synchronization between the stimulus presentation and the appearance of the trigger on the recorded EEG.

### 2.5 EEG data processing and analysis

#### 2.5.1. Data pre-processing

EEG preprocessing was performed in EEGlab/Matlab R2022b, ERPlab (Delorme & Makeig, 2004; Lopez-Calderon & Luck, 2014) and later replicated with Python/MNE (Gramfort et al., 2013).

Preprocessing steps used Independent Component Analysis (ICA) and were performed according to the recommended Makoto Miyakoshi EEGLab-pipeline (*Makoto’s preprocessing pipeline. (n.d.). Retrieved April 23, 2020, from https://sccn.ucsd.edu/wiki/Makoto’s_preprocessing_pipeline*) for each subject’s dataset individually. In more details, two steps were conducted :

1. To compute ICA weights, continuous raw EEG data were first filtered using a 30Hz-low-pass filter, a 1Hz high-pass filter (both 2^nd^order Butterworth filters at 6 dB/octave), and a 50Hz notch filter (Parks-McClellan filter). Data were then cleaned using the clean_rawdata EEGlab plug-in (Artifact Subspace Reconstruction, variance threshold 20). Then a full-rank ICA was performed using the Infomax algorithm.
2. To apply ICA weights, continuous raw EEG data were again low-pass filtered at 30 Hz, high-pass filtered at 0.1 Hz with a 50Hz notch filter. Bad electrodes (defined by an amplitude standard deviation < 2 or > 100μV) were interpolated with the trimOutlier EEGlab plug-in. Data were then cleaned with the clean_rawdata plug-in. ICA weights obtained at the first step were then applied to this EEG dataset. The IClabel EEGlab plugin was used to label independent components (among 7 labels : brain, muscle, eye, heart, line noise, channel noise and other), and components reflecting eye artifacts were removed. EEG data were then re-referenced to the average of all electrodes (and Cz electrode was added). Finally, continuous EEG data were epoched using a time-interval [-100; +600ms] relative to the auditory stimulus onset, and baseline-corrected.

#### 2.5.2. Event-related potentials analysis

Event-related potentials for the N= 24 participants were obtained by averaging epochs for each of the 6 deviant conditions (pitch conditions: rising, falling, flat; timbre conditions: smile, rough, neutral). Grand-averages were computed by averaging participant-averages across participants using the ERPlab software (Lopez-Calderon & Luck, 2014).

First, to explore the differences between the deviant own-names conditions without any a priori about the kind of differences in terms of topography or time course, we conducted cluster-based permutation tests on all 64 EEG sensors using the EEGLab FieldTrip plugin (Oostenveld et al., 2011) within a latency range of [0;600ms]. Separate cluster-based permutation tests were done for each prosody contrast in both the pitch (Rise versus Fall, Rise versus Flat, Flat versus Fall) and timbre conditions (Smile versus Rough, Smile versus Neutral and Neutral versus Rough). Cluster permutation tests used the Monte-Carlo method with 1000 permutations.

To better characterize electrophysiological responses within the spatio-temporal clusters identified with permutation tests, we then performed an additional parametric analysis using the ERPlab software (Lopez-Calderon & Luck, 2014). We calculated the peak amplitude over 3 ms and the area under curve (AUC) in each condition in the latency range corresponding to the significant clusters, and on electrodes representative of its spatial extent. We then tested for statistical differences across conditions, within subjects, using repeated measures ANOVA in the Jasp software (JASP Team (2022), version 0.16.3).

#### 2.5.3 Source localization

Finally, to better understand which brain areas were differently activated by deviant own-name conditions, we performed a source localization analysis when the cluster-based permutation tests displayed significant differences between conditions. First, we computed a mean head model (Gramfort et al., 2010) for all participants to perform source localizations with Brainstorm (Tadel et al., 2011). A noise covariance matrix was computed for each participant by taking the 100ms baseline period of each trial. We computed, for each subject, one sensor-level average per condition (rising, falling and neutral conditions; and smiling, rough and neutral conditions). We then estimated sources for each average according to the minimum norm imaging (MNI) method on the whole time interval [-100; 600ms], using two methods of standardization: dynamical Statistical Parametric Mapping (dSPM; (Dale et al., 2000)) and standardized Low resolution brain Electromagnetic Tomography (sLORETA) (Pascual-Marqui, 2002).

Source cortical maps were then compared with permutation paired t-test (with Flase Discovery Rate – FDR-correction) between deviant types, in the time-window and in ROIs identified as significant in the sensor-level cluster-based permutation test. Some ROIs were also independently investigated, because they have been described as relevant for the generation of late auditory evoked potentials in previous literature (Halgren et al., 1998; Perrin et al., 2005): posterior cingulate gyri, supramarginal gyri, superior temporal sulci, ventrolateral and dorsolateral and medial prefrontal cortices.

## 3. Results

### 3.1 Behavioral results

Participants were asked to covertly count the number of own-name deviants in both sequences. Participants performed relatively accurately, with a mean count of 233 / 240 true deviants. There was no significant difference in participants’ performance between paradigms (pitch paradigm: M = 232.3/240; timbre paradigm: M = 233.5/240 own-name deviants, t(16) = 0.12, p = 0.91; 7 participants had missing values due to technical problems or losing count).

### 3.2 ERP grand-average in both paradigms

ERP grand-averages for both pitch and timbre paradigm are shown in Figure 1. We observed a clear N100 response for both standard tones and deviant own-names. We also observed a wide positive deflection for all deviant own-names (and not for standard tones), between 220 and 350ms, with a maximal amplitude at the level of centro-parietal sensors (Figure 1). Because of its spatial distribution (centro-parietal), its latency and the kind of auditory paradigm that we used (oddball auditory paradigm with a covert counting task of deviant stimuli), in the rest of this manuscript, we therefore consider this positive deflection as a “P300 response” to deviant own-names.

**Figure 1:**
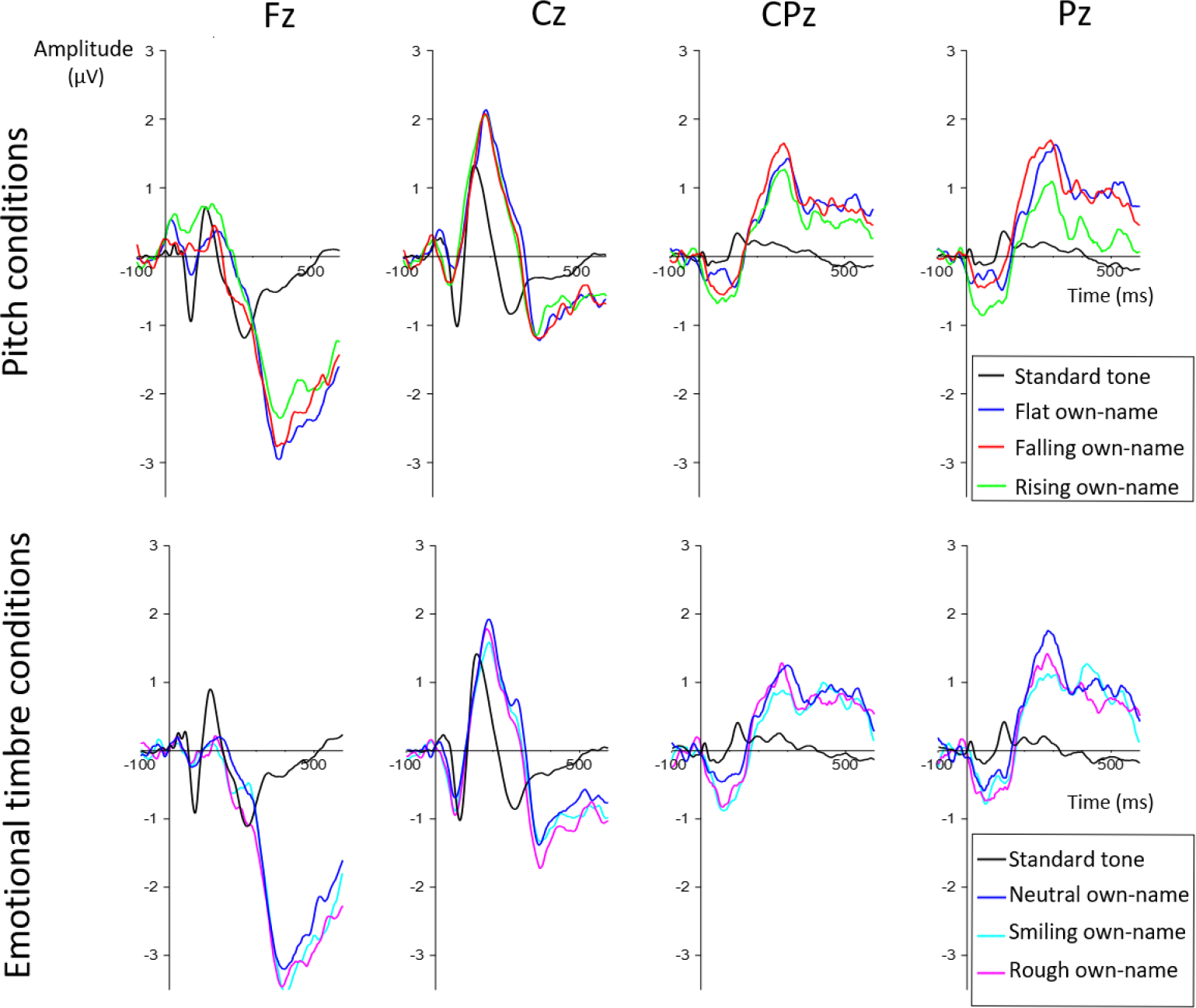
Grand-average ERPs in the pitch (top) and emotional timbre paradigms (bottom), for a selection of frontal to parietal midline sensors (left to right). As expected with oddball paradigms, frequent standard sounds elicited higher N100 responses than rarer deviants (seen here e.g. on Fz and Cz), while deviant sounds generated a clear P300 response on central parietal sensors. Rising pitch contours elicited statistically smaller P300 amplitudes than falling pitch contours (top-right), while emotional timbre conditions did not have any observable impact on P300 amplitude (bottom-right).

In the following, we analyze the pitch and timbre paradigms separately.

### 3.3 Pitch paradigm

Topographies of the grand-average ERP in all three pitch conditions (rising, falling and flat) are shown in Figure S1.

We first ran three separate cluster-based permutation tests to compare rising and falling pitch, rising and flat pitch, and falling and flat pitch in the [0-600ms] time-window. The comparison of rising vs falling pitch was significant (p<0.05) in a parieto-occipital cluster between [180;320ms], more pronounced on Pz (Figure 2-top). There was no significant difference between rising and flat, and between falling and flat.

**Figure 2:**
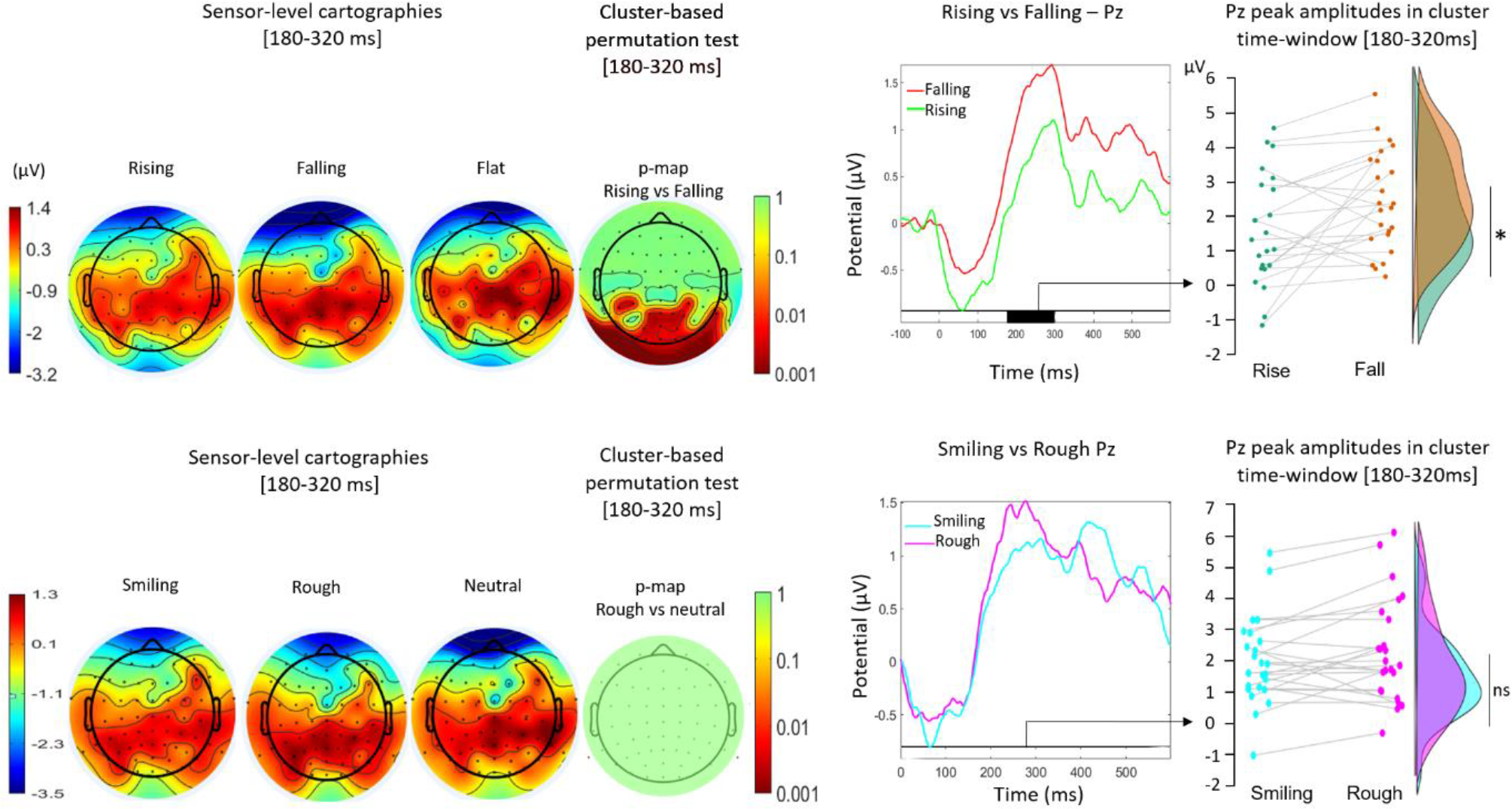
Sensors-level cartographies, cluster-based permutation test and parametric results in pitch (top) and emotional timbre (bottom) conditions. Top, from left to right: Sensors-level cartographies in the most discriminative time-window for the pitch paradigm [180-320 ms]; cluster-based permutation t-map in the same time window (t-values are represented with a colored scale ranging from 1 to 0.001 or from green to red); Pz sensor data comparing rising and falling deviants: raincloud plot comparing Pz peak amplitudes in [180-320 ms]. Rising prosodies elicited lower peak amplitudes than falling prosodies (p=0.02). Bottom, from left to right: Sensor-level cartographies, cluster-based permutation t-map (t-values are represented with a colored scale ranging from 1 to 0.001 or from green to red), Pz sensor data and raincloud plot comparing smile and rough deviants in the [180-320 ms] time-window. Contrary to pitch deviants, there was no significant difference between emotional timbre conditions.

To explore this parietal cluster in more details, we then averaged ERP data for rising and falling-pitch across a ROI comprising the 17 significant channels of the cluster (P7, P5, P3, P1, Pz, P2, P4, P6, P8, PO7, PO3, POz, PO4, PO8, O1, Oz, O2), and compared them in the [180-320ms] time-window using repeated-measure ANOVAs. In this cluster and in this time window, falling prosodies elicited both higher (peak amplitude: 1.7 versus 1.1 μV respectively, F(1,23) = 6.6, p = 0.017) and wider (AUC: 0.19 versus 0.15 μV/ms, F(1,23) = 5, p = 0.036) responses than rising prosodies (Figure 2).

**Figure S1:**
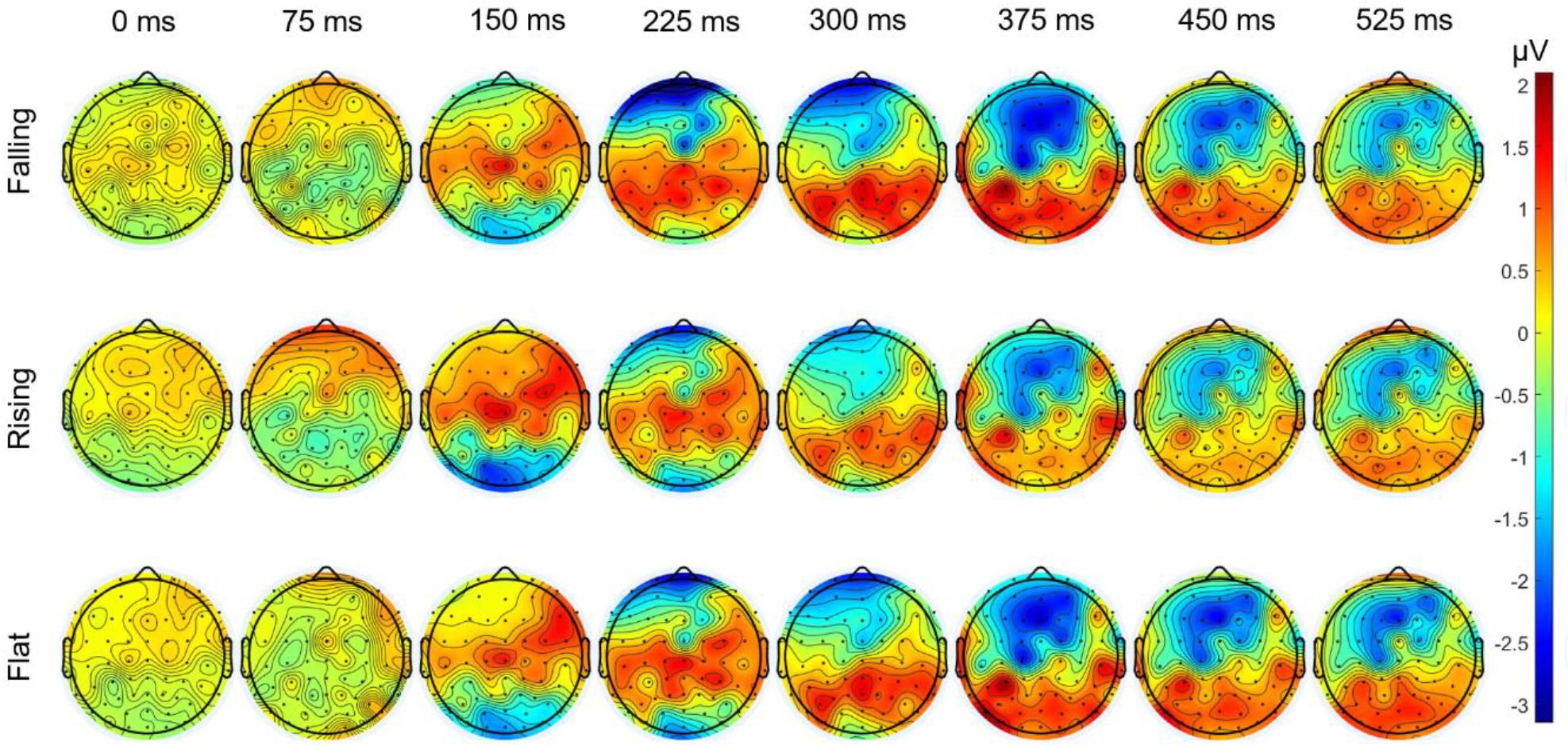
Grand-average ERP cartographies in pitch paradigm (between 0 to 525 ms post stimuli). Each trial in each pitch condition (rising, falling, flat) was averaged for all participants and instantaneous amplitudes are displayed on scalp maps.

Finally, we ran source analysis in the same time window, and checked for significant differences in source activations between rising and falling conditions. Falling deviants elicited significant lower activities than rising deviants in the left posterior cingulate gyrus (PCG) in both source localizations methods (between [180;247ms] with dSPM and between [225;246ms] with sLORETA).

Conversely, falling deviants elicited significant higher activities than rising deviants in the left superior frontal gyrus (SFG) with sLORETA (between [200;223] and [244;261ms]), although this difference was not reproduced with dSPM (Figure 3). Comparisons of sources in the supramarginal, temporal and ventrolateral prefrontal areas did not reveal significant differences between conditions.

**Figure 3:**
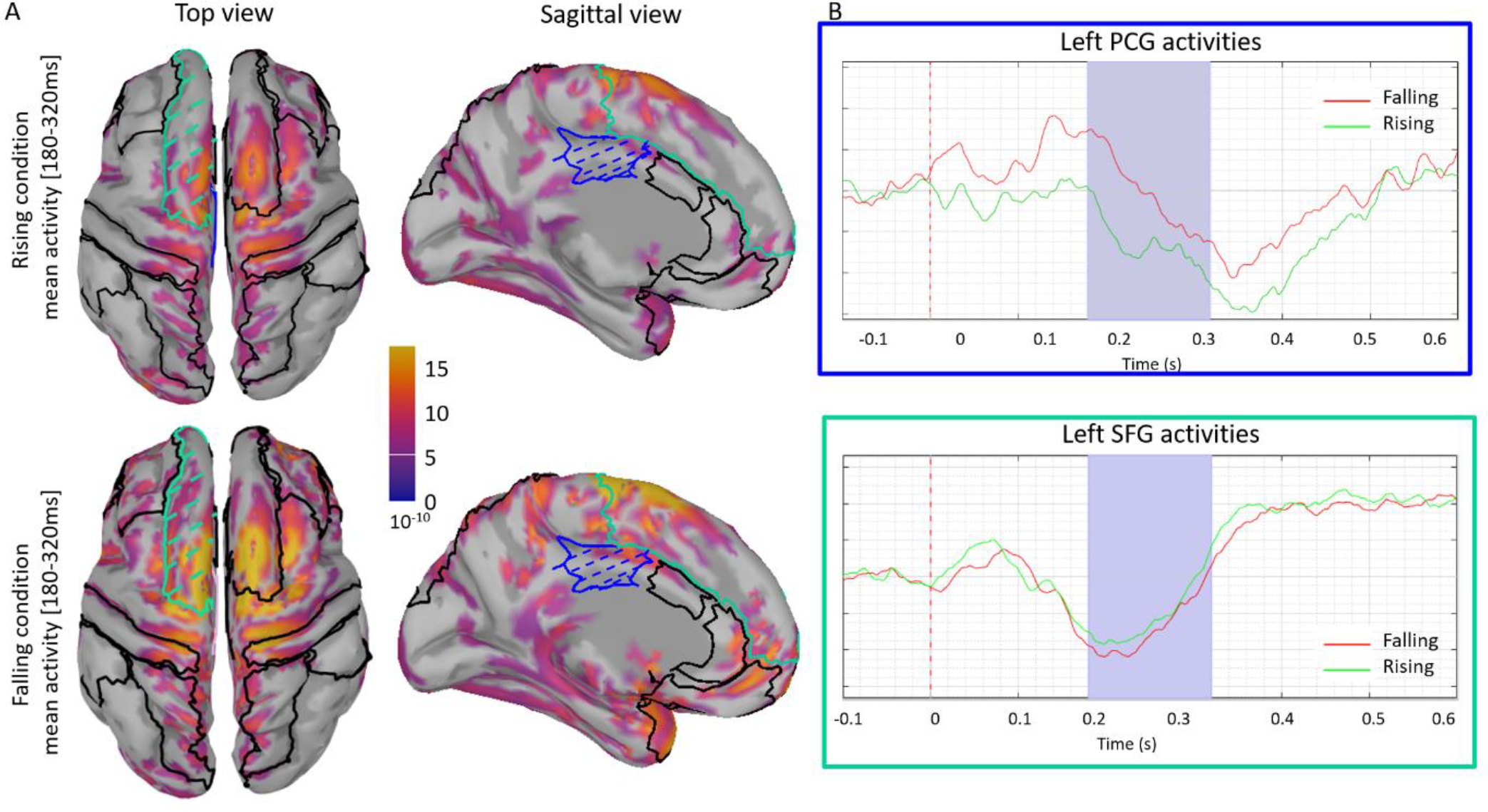
Source localization results in the pitch paradigm. A. Source localizations performed using sLORETA for the grand average of rising (up) and falling (bottom) conditions (top and sagittal views from left to right), in the [180-320ms] time window. Regions of interest (ROIs) included in the statistical comparison are circled in dark. The left posterior cingulate gyrus (PCG, hatched in blue) and the left superior frontal gyrus (SFG, hatched in green) displayed significant differences in the permutation t-test. B. sLORETA sources activities in (top) the left PCG and (bottom) the left SFG, for falling (red) and rising pitch (green). Source activities in these two ROIs differed between rising and falling conditions in the [180-320ms] time-window, highlighted in light purple.

#### Timbre paradigm

Topographies of the grand-average ERP in all three timbre conditions (smile, rough and neutral) are shown in Figure S2.

As for pitch, we first ran 3 separate cluster-based permutation tests to compare smiling and rough timbre, smiling and neutral timbre, and rough and neutral timbre in the [0-600ms] time-window. No significant difference was obtained.

To verify whether a less conservative analysis strategy would reveal weaker differences between timbre conditions, we then reduced the time-window of analysis of the cluster permutation test to either [200;400ms], [300;600ms] and [400;600ms]. Again, no significant difference was obtained.

Even so, we further explored the same time-window [180;320] identified as significant in the pitch paradigm. Cluster-based permutation tests between [180;320ms] did not reveal any significant results, nor did parametric analysis on Pz between the smiling and the rough conditions (Pz peak amplitudes: 1.9 versus 2.2 μV respectively, F(1,23) = 3.5, p = 0.07) (Figure 2).

**Figure S2:**
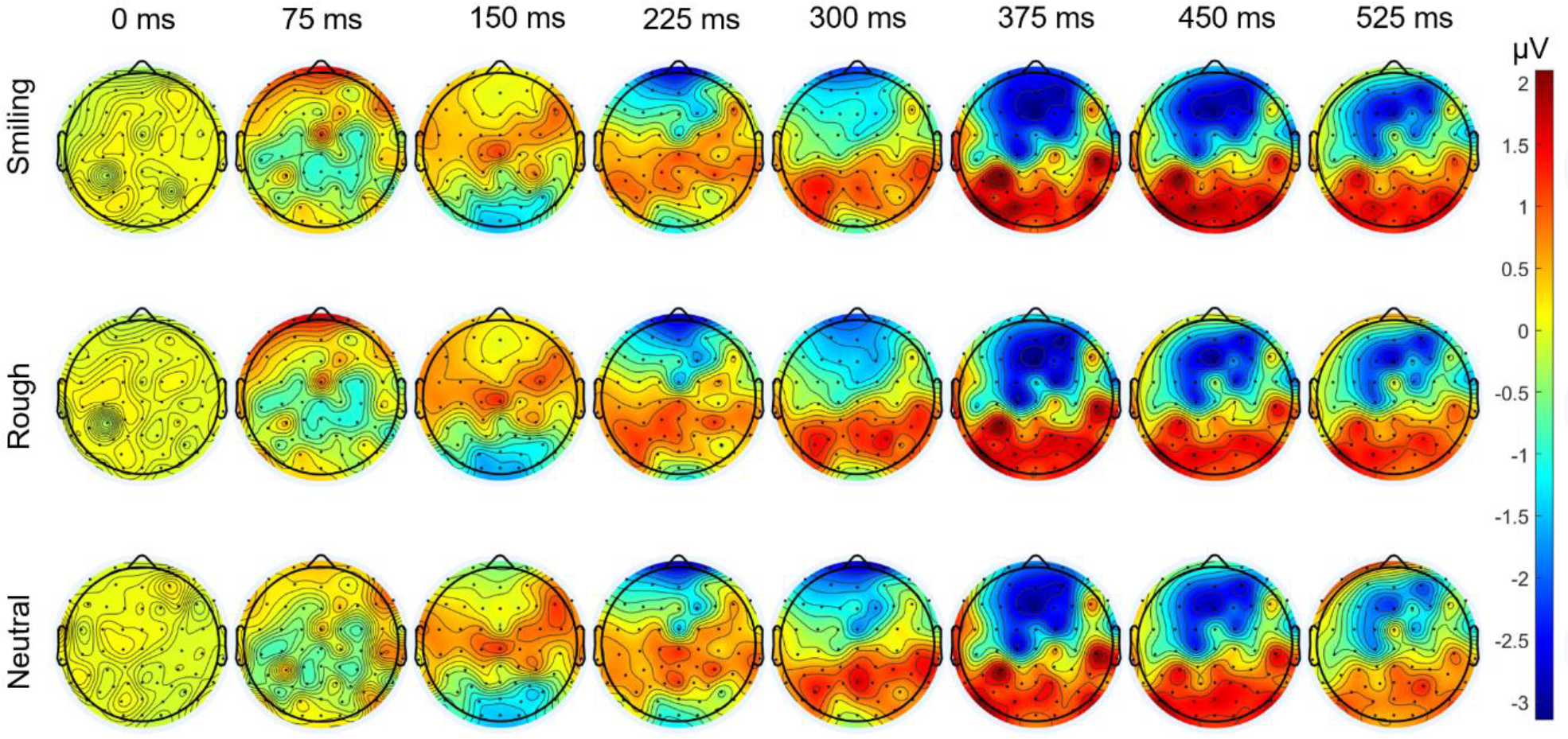
Grand-average ERP cartographies in emotional timbre paradigm (between 0 to 525 ms post stimuli). Each trial in each emotional timbre condition (smiling, rough, neutral) was averaged for all participants and instantaneous amplitudes are displayed on scalp maps.

## 4. Discussion

Because little is known about the electrophysiological impact of pitch or timbre modulations on a listener’s attention to speech stimuli when these variations are task-irrelevant, we adapted an auditory oddball paradigm routinely used in ICU to evaluate consciousness in DOC patients (the “own-name” paradigm) and tested whether systematic digital transformations of pitch and timbre influenced P300 responses in a group of N=24 healthy participants. While rising or falling pitch prosodies significantly modulated the participants P300 response (with falling pitch eliciting higher and wider responses in parieto-occipital areas), we found no such difference when comparing timbre conditions (smiling, rough or neutral).

The fact that the pitch contour of a target stimuli should influence the recruitment of task-driven attention in an oddball paradigm confirms, if at all needed, that dynamic variations of pitch are an essential element of prosodic processing (Schirmer et al., 2001; Tang et al., 2017) and that these variations are processed early and consequentially in the subsequent cognitive processing of speech and language.

This result rejoins a wealth of studies demonstrating that pitch and, more generally, prosodic variations in auditory targets can modulate mid-latency ERP components such as the MMN (Charpentier et al., 2018), P200 (Paulmann et al., 2013) or P300 (Nussbaum et al., 2022), in tasks often linked to the explicit evaluation of their emotion or semantics. What the present study adds, first, is finer control on what exact variation of pitch and timbre is applied to the stimuli and, second, the demonstration that these variations are processed and influence the recruitment of attention even in the absence of explicit evaluation objectives.

More specifically here, we found that own-name deviants with falling pitch contours elicited larger and wider P300 responses than deviants with rising pitch contours when the task was unrelated to the prosody of the target sounds. Utterances with rising pitch are typically regarded as more salient and more alerting: for instance, rising pitch is used to mark questions (Ponsot et al., 2018), surprise (Goupil et al., 2021; Lai, 2009) and to signal speaker unreliability (Goupil et al., 2021); in Goupil et al. (2021), words pronounced with rising pitch contours were associated with better accuracy and faster reaction times in a verbal working memory task. Our pattern of results, where falling and flat pitch preserved relatively large P300 responses while rising pitch decreased their amplitude, therefore suggests that rising pitch contours did not modulate the saliency of the target stimulus but, on the contrary, interfered with the participant’s attention in the counting task. Such an interpretation would be consistent by previous literature demonstrating similar effects where interfering auditory (e.g. infant cries) or visual stimuli decreased the participant’s attention in an unrelated task, resulting in smaller P200 (Dudek et al., 2016) or P300 amplitudes (Jocoy et al., 1998; Wilson et al., 2012). Here, the attention-grabbing properties of rising pitch stimuli would compete with task-related attention towards counting the occurrence of own-name regardless of their prosody, and in turn down-modulate the P300 response linked to the task. In our study we did not measure counting accuracy separately for rising and falling deviants, and so cannot confirm whether this electrophysiological effect had any behavioral consequences. It could therefore be interesting to extend the same paradigm with a button-press detection task, and look for increased error rate or slower reaction times in rising versus falling trials.

The differential brain processing of falling and rising pitch prosodies also appeared in source activations within the left superior frontal gyrus and the left posterior cingulate gyrus. These source activations have to be interpreted with caution because participants did not underwent morphological MRI nor measurement of electrode positions. However, these activation are consistent with previous studies. Indeed, cingulate gyri have been implicated in attentional processes (Posner & Dehaene, 1994) and, along with frontal cortices, in the generation of P300 responses (Halgren et al., 1998). The differential source activations seen here are thus consistent with an influence of pitch-deviant stimuli on the attention given to the counting-task and its related P300 response. Alternatively, the modulation of left frontal activity seen in our participants may also reflect involvement in prosodic decoding. While prosodic processing is traditionally associated with right temporo-frontal networks (Schirmer & Kotz, 2006), previous literature has indeed demonstrated the possible wider activation of bilateral inferior frontal, prefrontal and caudate nucleus areas during effortful prosodic speech listening (Kotz et al., 2003). It is therefore possible that left frontal activation in pitch contours conditions reflects the specific involvement of cognitive resources to decode prosodic information, the same resources that may compete with task-relevant resources for counting own-name deviants.

In contrast to pitch contours, our study shows a striking lack of evidence for any effect of timbre modulations. The fact that own-name deviants were transformed to sound more smiling or rougher than normal had no observable consequence on the participants’ P300 responses.

One methodological reason why one would not record evidence of any effect of timbre would be that the two timbre manipulations (smile and rough) used in this study were somehow less acoustically effective than the two pitch manipulations. While we did not measure ratings of e.g. emotionality of the four types of stimuli in the same participants, we find this explanation implausible because both algorithms have been associated with very large effects in other studies measuring explicit evaluations of emotional valence and arousal (smiling, more positive and more aroused; rough, more negative and more aroused – (Arias et al., 2018; Bedoya et al., 2021; Ollivier et al., 2019).

Another reason for the absence of a clear evidence of modulation of the P300 characteristics in the timbre paradigm could be the implication of deeper cerebral areas which could have been missed with surface recordings. For instance, the rough quality of a voice has been associated not only to frontal and auditory activities but also to the synchronization of limbic areas (and notably hippocampus and insula) in intracranial recordings (Arnal et al., 2019). Surface recordings may not be sufficiently sensitive to detect such effects. However, we again find this explanation implausible, because pitch contours effects in the first paradigm were associated with clear source differences in cortical areas involved in P300 generation, such as the PCG, and nothing would have prevented registering similar effects in timbre comparisons.

In short, our results suggest that timbre modulations (smiling/rough) do not have the same capacity as pitch contours (rising / falling) to capture bottom-up attention, and/or interfere with task-related attention towards counting the occurrence of own-name regardless of their prosody.

In our view, the main difference between both types of stimuli is that smile and rough variations primarily carry emotional meaning, while pitch contours warrant a wider variety of cognitive evaluations, including emotions but also social attitudes or linguistic prosody (Wichmann, 2000). There is a long-ranging debate regarding the automaticity of emotional processing, i.e. whether such stimuli can be processed without requiring attention or whether their processing competes with general-domain attention resources (Pourtois et al., 2013). In some studies, brain regions responding to e.g. emotional faces were found to do so even when attention is guided away from the stimuli, e.g. with a spatial visual task (Vuilleumier et al., 2001); in others however, amygdala responses to angry faces or voices were reduced when presented under high attentional load (Mothes-Lasch et al., 2011; Pessoa et al., 2002). In our participants, multiple attention gain control systems (Pourtois et al., 2013) likely operated in parallel: bottom-up processes attending to time-varying pitch cues, to emotional signatures in vocal timbre, and top-down processes attending to features allowing to distinguish vocal own-name deviants from non-vocal standard sounds. It appears possible that these three systems are mediated by distinct mechanisms that do not necessarily compete on the same neural resources: for instance, the emotional timbre processing of smile and roughness may only weakly compete with the cognitive/attention resources required by our task; alternatively, the sensory resources involved in pitch contour processing may overlap more strongly with those required to count own-name deviants (e.g. auditory cortical areas linked to voice recognition) than sensory resources involved in emotional timbre processing, which are maybe mediated by distinct neural mechanisms in amygdala and interconnected prefrontal area.

In this view, it therefore remains possible that effects of emotional timbre could be observed if participants engaged in a task with higher-attentional load (e.g. detecting own-name with a superposed brief tonal target) or that recruits slightly different sensory cues that more closely overlapped timbre cues (e.g. detecting own-name pronounced by a specific speaker). In addition, it is also possible that bottom-up emotional timbre processing did compete with the processing of acoustic features allowing to detect own-name deviants, but at a finer temporal or spatial scale than we could observe here. For instance, using intracranial recordings, Pourtois et al. (2010) recorded both early (140–290ms) amygdala responses to threatening faces that arose independently of attentional focus and, at a later time (starting at 700ms) an attention-dependent response - see also (Luo et al., 2010). It would be therefore interesting to investigate the effect of smile and rough modulations in the own-name task using intracranial recordings or magneto-encephalography (Pizzo et al., 2019).

Finally, because own-name oddball paradigm investigated in this study is widely used to assess DOC patients (André-Obadia et al., 2018), our present results also have clinical implications. In typical patient evaluations, the acoustic characteristics of the own-name stimuli are left relatively uncontrolled, e.g. because they are recorded by staff or family members. Our results suggest, first, that normalizing the pitch contour of deviant sounds to e.g. a falling prosody could facilitate their registering by a patient’s attentional system and improve the signal-to-noise ratio for detecting otherwise typically weak P300 signatures in the patients (Pruvost-Robieux, André-Obadia, et al., 2022). Second, it is also possible that observing a modulation of P300 by prosodic features such as pitch contours is in itself a marker of high-functional cortical processing (i.e. capable of sophisticated sensory processing and/or flexible attentional allocation), which can be indicative of better prognosis upon awakening (see e.g. (Bekinschtein et al., 2009)).

In a previous retrospective study (Pruvost-Robieux, André-Obadia, et al., 2022), we tested the impact of the (arbitrary) pitch contours used when recording own-name stimuli for testing P300 responses in N=251 DOC patients; here we used a similar paradigm on healthy participants. While both studies confirm a general role of pitch contours in modulating own-name P300 responses, they differed in their specific conclusions: in DOC patients, we found that stimuli recorded with a rising-pitch intonation were associated with shorter P300 latencies (but we couldn’t test amplitudes); here, we found rising-pitch intonations cause smaller P300 amplitudes (and found no effect on latency). In both studies however, the block-structure of stimuli differed: while in the present study rising and falling-pitch stimuli alternated within a block, DOC patients were tested with only one type of stimulus in a single block (e.g. all own-name deviants had rising pitch). It is therefore possible that, in that latter case, the repetition of attention-grabbing rising-pitch stimuli did not so much compete with task-related attention, but rather elevated the general level of attention, leading to beneficial side-effects on the main task. We can assume that the effects found here in healthy participants presuppose a fully functional attentional system, able to flexibly moderate endogenous and exogenous capture, and that the same mechanisms lead to different cognitive outcomes in patients with disorders of attentional systems. Further prospective studies in DOC patients will therefore be required to clarify the interplay between consciousness, and the modulation of top-down attention by prosodic and emotional cues.

## Abbreviations

DOC: (Disorders of consciousness)
ERPs: (Event related potentials)
ICU: (Intensive Care Units)
MMN: (mismatch negativity)
AUC: (Area under curve)
SFG: (Superior Frontal Gyrus)
PCG: (Posterior Cingulate Gyrus)
MNI: (minimum norm imaging)
dSPM: (dynamical Statistical Parametric Mapping)
sLORETA: (standardized Low resolution brain Electromagnetic Tomography)
ROI: (Region Of Interest)
RMS: (Root Mean Square)

## Data availability

*The data underlying this article will be shared on reasonable request to the corresponding author*.

## Funding

This work was supported by European Research Council [StG CREAM 335536], Agence Nationale de la Recherche [Sounds4Coma ANR-19-CE19, SEPIA ANR-19-CE37], Fondation pour l’Audition [FPA RD-2018–2] (to JJA) and “Sauver la vie 2020” grant (to EPR).

Funders had no involvement in the study.

## Acknowledgements

This study was conducted with the support of the INSEAD-Sorbonne Université Multidisciplinary Center for Behavioral Science.

## Declaration of Competing Interest

None of the authors have potential conflicts of interest to be disclosed.

## Notes

### Competing Interest Statement

The authors have declared no competing interest.

